# Encoding of social exploration by neural ensembles in the insular cortex

**DOI:** 10.1101/843490

**Authors:** Isamu Miura, Masaaki Sato, Nobuo Kunori, Eric T.N. Overton, Junichi Nakai, Takakazu Kawamata, Nobuhiro Nakai, Toru Takumi

## Abstract

The insular cortex participates in diverse complex brain functions including sociality, yet little is known about their cellular bases. Using microendoscopic calcium imaging of the agranular insular cortex (AI) in mice interacting with freely-moving and restrained social targets, we identified two subsets of AI neurons –a larger fraction of Social-ON cells and a smaller fraction of Social-OFF cells– that change their activity in opposite directions during social exploration. Social-ON cells included those that represented social investigation independent of location and consisted of multiple subsets, each of which were preferentially active during exploration under particular behavioral state or with a particular target of physical contact. These results uncover a previously unknown function of AI neurons in encoding conjunctive information on social behavior and suggest that AI may act to monitor the ongoing status of social exploration while an animal interacts with unfamiliar conspecifics.

## Introduction

Social animals communicate and interact with conspecifics both cooperatively and competitively for reproduction and survival. Social interaction between two individuals typically begins with an appetitive phase that reflects interest in peers, such as approaching and investigation by touching and sniffing, followed by a consummatory phase that express a repertoire of goal-directed social behaviors, such as aggression, mating or parenting, according to age, gender and familiarity of the conspecific (Anderson, 2016; Chen and Hong, 2018). Recent advances in social neuroscience in rodents have revealed critical centers for social behavioral control such as the medial preoptic area of the hypothalamus (MOPA) (Wu et al., 2014; McHenry et al., 2017), the posterodorsal and posteroventral medial amygdala (MeApd) (Hong et al., 2014; Li et al., 2017), the ventrolateral subdivision of the ventromedial hypothalamus (VMHvl) (Lin et al., 2011; Remedios et al, 2017) and the medial prefrontal cortex (mPFC) (Wang et al., 2011; Yizhar et al., 2011; Murugan et al., 2017; Liang et al., 2018; Kingsbury et al., 2019). However, functional characterization of additional network nodes is essential for the full understanding of neural circuit mechanisms underlying complex social behavior.

The insular cortex (IC), which lies deep within the lateral sulcus in humans and on the lateral aspect of the neocortex in rodents, is involved in a wide variety of functions including multisensory integration, emotion, risk prediction, decision making, salience and valence coding, empathy and self-awareness (Craig, 2009; Gogolla et al., 2014; Uddin, 2014; Gogolla, 2017; Rogerss-Carter et al., 2018). It forms an anatomical hub with reciprocal connections to sensory, emotional, motivational and cognitive systems including the sensory and frontal cortices, amygdala, thalamus and nucleus accumbens as well as with neuromodulatory inputs (Allen et al., 1991; Shi and Cassell, 1998; Jasmin et al., 2003; Kobayashi, 2011). Functional magnetic resonance imaging studies in humans suggest that IC serves as the core of a “salience network” that acts to detect novel and behaviorally relevant stimuli (Seeley et al., 2007; Uddin, 2014). Furthermore, hypoactivity and dysfunctional connectivity of IC are hallmarks of individuals with autism spectrum disorder (ASD) (Uddin and Menon, 2009; Di Martino et al., 2009). These findings imply that IC constitutes a key node for the social brain network and plays a potential role in social recognition. In this study, we sought to examine how neurons in IC encode information regarding social behavior by visualizing neuronal dynamics using microendoscopic calcium imaging in freely-moving, socially-interacting mice. Our study particularly focused on the agranular insular cortex (AI), the most anterior part of the IC that lacks the granular layer 4 and has denser limbic connections than the rest of IC (Allen et al., 1991; Kobayashi, 2011).

## Results

### Multiple subsets of insular cells encode social exploration behavior

To elucidate how AI neurons encode direct social interaction with unfamiliar conspecifics, we first conducted a home-cage (HC) experiment (Silverman et al., 2010), in which a male mouse was allowed to explore a novel object placed in its HC for the first 4 min, followed by another 4 min of exploration of a freely-moving male stranger mouse that was introduced to replace the object (Fig. 1A). While subject mice displayed only occasional nasal contacts with the novel object during control sessions, they showed highly frequent nasal contacts with the stranger mice during interaction sessions (Fig. 1B-C). The subject mice spent 30.5% of the total time on social interaction, defined by their nasal contact to the stranger mice (Fig. 1D). Furthermore, 48.6%, 45.1% and 6.3% of the social interaction period were spent in contact with the nose, body and anogenital area of the stranger mice (Fig. 1D and 1E). Besides contact initiated by the subject mice, the stranger mice also contacted the subject mice for 6.2% of the total time. During periods where the subject was not interacting with the stranger, the subject mice touched the wall of their HCs for 0.6% of the total time. The subject mice remained stationary for 70.0% of the total time, and the fraction of stationary periods increased to 82.3% when only the social interaction periods were considered.

**Figure 1.**
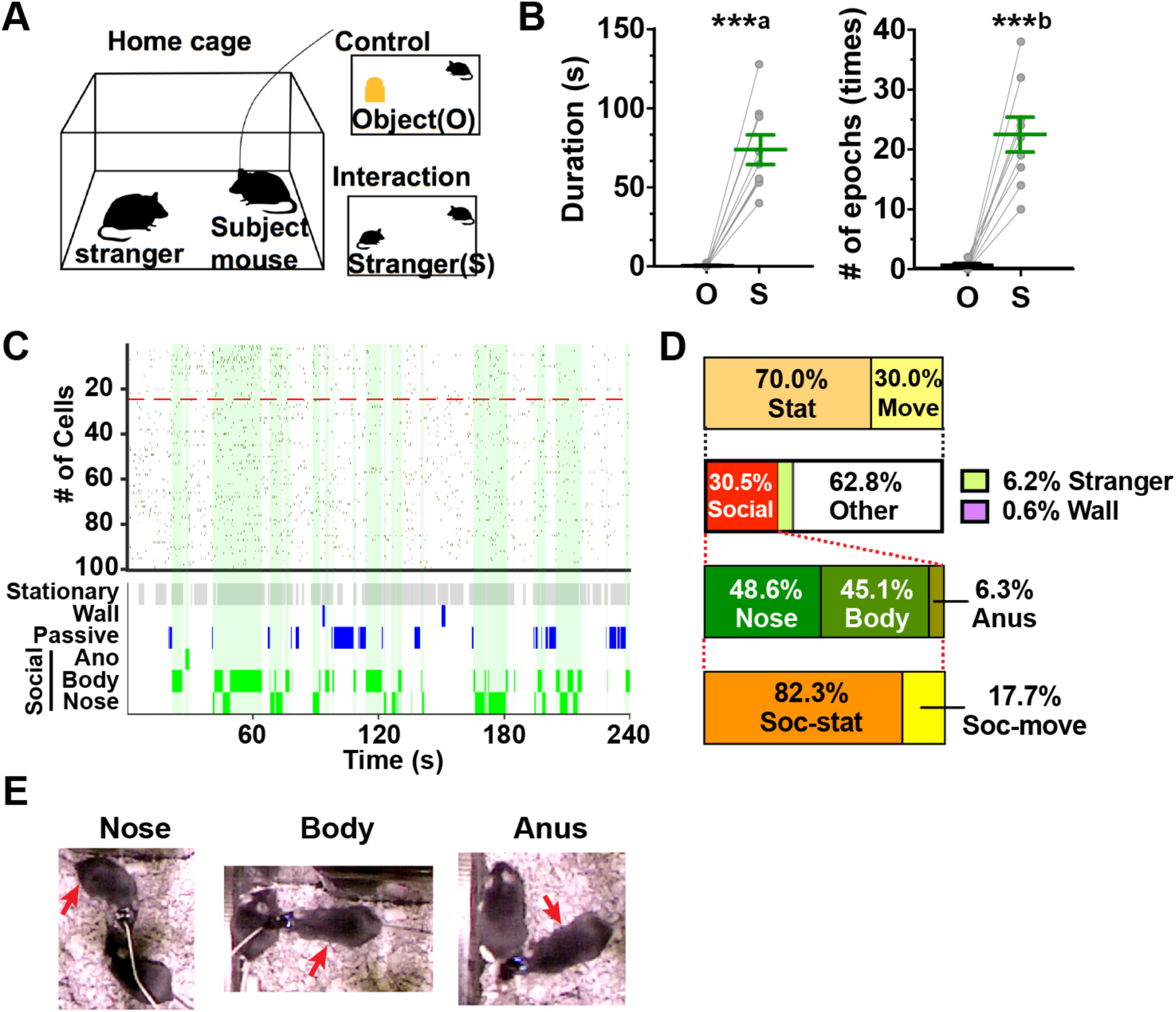
Behavior during HC experiments. (A) Experimental design of HC experiment. A subject mouse was allowed to explore a novel object in its HC for 4 min in a control session (top right), followed by an additional 4 min of an interaction session with a stranger mouse (bottom right). (B) Total duration (left) and the number of nasal contact epoches (right) with a novel object (O) during control sessions and with a stranger mouse (S) during interaction sessions. ***a, P < 0.0001 vs. O, t_(8)_ = 7.902, unpaired t-test. ***b, P < 0.0001 vs. O, t_(8)_ = 7.348, unpaired t-test. (C) A representative raster plot showing Ca^2+^ events of a population of AI neurons imaged in a single experiment (top, n = 100 neurons). The epochs of social interaction are indicated in green. Social-ON cells are sorted above the red dashed line. Behavior of the mouse is shown in the bottom panel, in which non-moving periods (stationary), contact with the wall (wall), passive touch by the stranger mouse (passive) and contact with the anogenital area (ano), body and nose of the stranger are presented from top to bottom. (D) Characterization of the behavior of subject mice in HC experiments. The fractions of time spent moving (move) and not moving (stat) and those for social interaction (social), passive touch by the stranger (stranger), contact with the wall (wall) and otherwise in entire sessions are shown in the top and upper middle bars, respectively. The bars presented in the lower middle and bottom indicate the fractions of time spent touching the nose, body and anogenital areas (anus) and those spent moving (Soc-move) and not moving (Soc-stat) during the periods of social interaction, respectively. (E) Sample video frames of contact with the nose, body and anus of the stranger. The subject mouse has a miniaturized microscope attached on its head as indicated by the red arrow.

We then sought AI neurons that exhibited social interaction-associated activity using microendoscopic calcium imaging in subject mice (Supplementary Fig. 1). We calculated cosine similarities between the vectors representing their binarized neuronal activity and those representing when the subject mouse interacted with the stranger mouse. In the analysis, we selected cells whose activity was significantly correlated or anti-correlated with the social interaction period but not with generic behavioral states such as either moving or stationary periods. Out of 482 cells from 9 mice, we identified 20.1% (“Social-ON cells”, 97 cells) and 3.1% (“Social-OFF cell”, 14 cells) of total cells whose activity was correlated and anti-correlated with social interaction period, respectively (Fig. 2A-B). The average event rate of Social-ON cells during periods of no social interaction (“non-social period”) was significantly lower than cells that were neither Social-ON nor Social-OFF cells (“non-social cells”) and substantially increased during the social interaction period compared to the non-social period. In contrast, the average event rate of Social-OFF cells during the non-social period was significantly higher than that of the non-social cells and markedly decreased during social interaction compared to the non-social period (Fig. 2C-D). As a result, activity of Social-ON cells and Social-OFF cells exhibited strong biases toward the social interaction period and the non-social period (Social-ON cells, 0.672 ± 0.165, n = 97 cells; Social-OFF cells, −0.850 ± 0.167, n = 14 cells), respectively (Fig. 2E). Within the microendoscopic field of view, Social-ON cells and Social-OFF cells did not appear to form segregated clusters, but rather coexisted in an intermingled manner (Fig. 2F). The nearest neighbor distances between Social-ON cells calculated using real and shuffled data did not differ significantly, indicating that Social-ON cells are distributed randomly within the AI (Fig. 2F).

**Figure 2.**
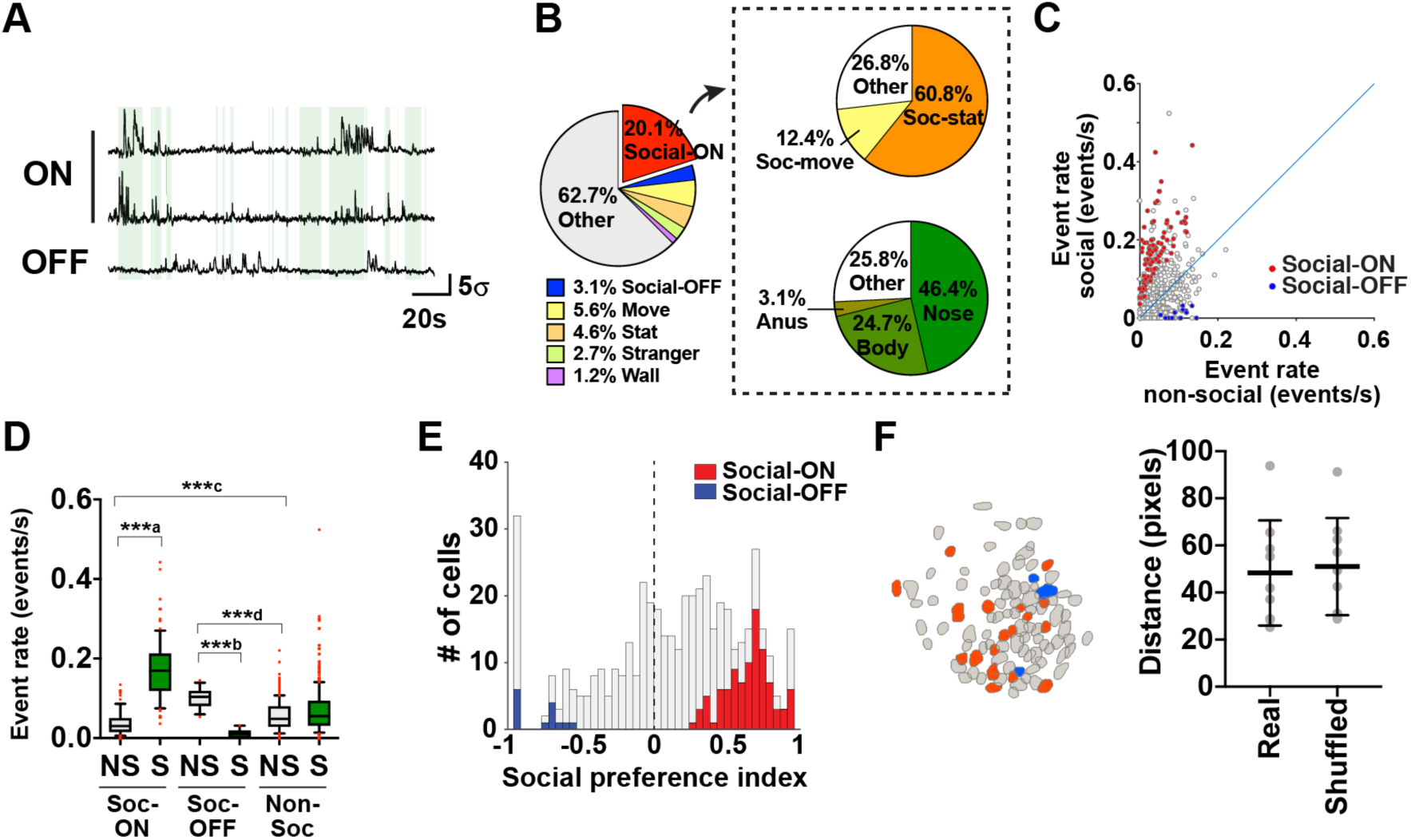
Social-ON cells and Social-OFF cells in AI. (A) GCaMP ΔF/F traces of two representative Social-ON cells and a Social-OFF cell. The epochs of social interaction are indicated in green. (B) A pie chart showing the fractions of each category of cells in the total cell population (left, n = 482 cells from 9 mice). The pie charts shown within the dashed rectangle demonstrate the fractions of each subcategory of cells in the Social-ON cell population. (C) A scatter plot of Ca^2+^ event rates of individual neurons during social interaction and non-social periods (n=482 cells from 9 mice). The blue line indicates equivalence. Social-ON cells (n = 97 cells) and Social-OFF cells (n = 14 cells) are shown in red and blue, respectively. (D) Box plots of Ca^2+^ event rates of Social-ON (Soc-ON, n = 97 cells), Social-OFF (Soc-OFF, n = 14 cells) and non-social cells (Non-soc, n = 371 cells) during non-social (NS) and social interaction (S) periods. Whiskers represent 10–90 percentile and red dots represent outliers. ***a, P < 0.0001, W_(97)_ = 4753, n = 97 cells; ***b, P = 0.0001, W_(14)_ = −105, n = 14 cells; Wilcoxon matched-pairs sign rank test; ***c, P < 0.0001, U_(97, 371)_ = 12375; n = 97 and 371 cells; ***d, P < 0.0001, U_(14, 371)_ = 834.5; n = 14 and 371 cells; Mann-Whitney test. (E) Distribution of social preference indices of individual neurons. The fractions of Social-ON cells, Social-OFF cells and non-social cells are shown in red, blue and gray in stacked bars. (F) Anatomical distribution of Social cells in a representative field of view (left, n = 100 cells). Social-ON cells (n = 18 cells) and Social-OFF cells (n = 3 cells) are shown in red and blue, respectively. Average nearest neighbor distances between Social-ON cells calculated using real (Real) and shuffled data (Shuffled) are shown in the right panel (P = 0.40, Real vs. Shuffled, t_(8)_ = 0.893, paired t- test, n = 9 mice).

In addition to these Social-ON and Social-OFF cells, we identified 5.6% (27 cells) and 4.8% (23 cells) of total cells as cells that exhibited activity significantly correlated with overall moving and stationary periods, respectively. These cell categories displayed higher event rates during their relevant behavioral state compared to the other state (termed “Move cells” and “Stat cells”, respectively), irrespective of whether they were socially interacting or not (Fig. 2B and Supplementary Fig. 2A–B). Interestingly, 60.8% (59 cells) and 12.4% (12 cells) of Social-ON cells also exhibited activity significantly correlated with the parts of social interaction periods during which the subject mouse was stationary and moving, respectively (termed “Social-stat cells” and “Social-move cells”; Fig. 2B and Supplementary Fig. 2C). These subcategories of cells demonstrated higher event rates during the relevant behavioral state only when the subject mouse was socially interacting (Supplementary Fig. 2D), indicating that subsets of insular Social-ON cells are preferentially activated while a subject mouse interacts with a stranger under a particular behavioral state. Finally, we examined if specific social contact by the subject mice also influenced the activity of Social-ON cells. We found that 46.4% (45 cells), 24.7% (24 cells) and 3.1% (3 cells) of Social-ON cells showed activity that was significantly correlated with the period during which they contacted with the nose, body and anogenital area of the stranger mice (Fig. 2B and Supplementary Fig. 2E–F), although the amount of time spent contacting with the anus was low (n = 4 out of 9 mice; range 0.7–7.3% of total time; Fig. 1D). In summary, these findings suggest that insular Social-ON cells do not merely respond to social interaction in general but rather represent conjunctive information regarding behavioral state and the target of contact during social interaction.

### Location-independent coding of social investigation by AI neurons

We next investigated activity of AI neurons in a linear-chamber (LC) apparatus, a modified version of the three-chamber test that can control spatial factors and physical contact (Fig. 3A; Lee et al., 2016). In these experiments, male subject’s preference for social stimuli was assessed by comparing the amount of time spent investigating two separate small chambers which contained either a male stranger mouse or a novel inanimate object. The investigation behavior was defined as poking of the subject mouse’s nose into holes of the acrylic wall of the small chambers. The transparency of the wall and its holes allowed transmission of olfactory, visual, auditory and limited tactile social cues to the subject mice, while preventing full physical contact of the subject mice with the target.

**Figure 3.**
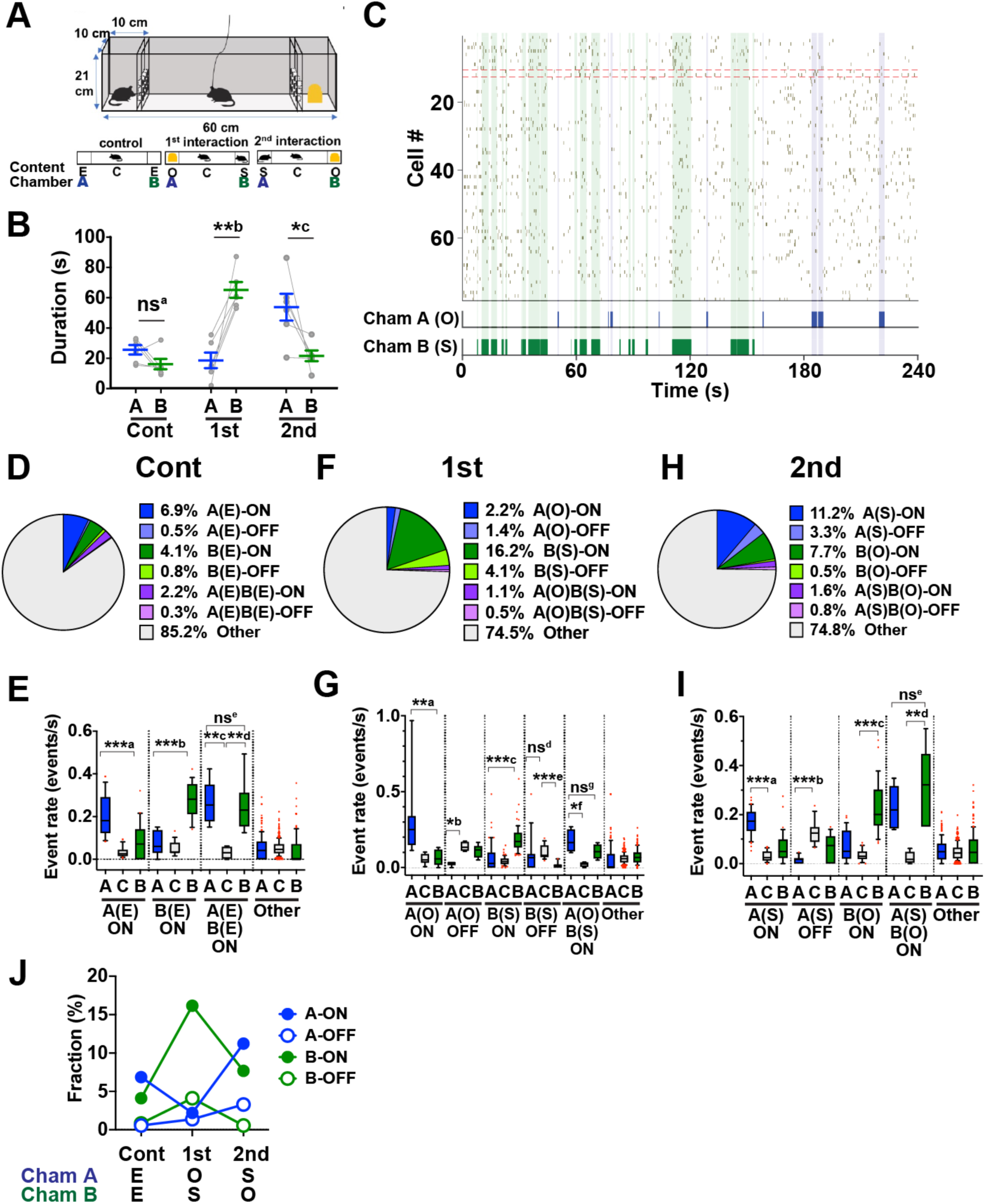
Behavior and AI neuron activity during LC experiments. (A) Experimental scheme. Mice were subjected to three 4 min-sessions in the LC (top) in the following order (bottom, from left to right); control session in which chambers A and B were empty (E), the first social interaction session in which chamber A and chamber B contained a novel object (O) and a stranger mouse (S), respectively, and the second social interaction session in which chamber A and chamber B contained a stranger mouse and a novel object, respectively. (B) Total duration of time spent investigating chamber A and chamber B during control (Cont), first interaction (1st) and second interaction (2nd) sessions. ns^a^, P = 0.12, t_(5)_ = 1.89; **^b^, P = 0.0022, t_(5)_ = 5.75; *^c^, P = 0.036, t_(5)_ = 0.036, paired t-test, n = 6 mice. (C) A representative raster plot showing Ca^2+^ events of a population of AI neurons (n = 78 cells) imaged in a single experiment during the 1st interaction session. B(S)-ON and B(S)-OFF cells are sorted above the red dashed lines. The epochs of nose-poking to chamber A with a novel object (Cham A (O)) and chamber B with a stranger mouse (Cham B (S)) are shown in the bottom panel and indicated by blue and green shades, respectively. (D) A pie chart showing the fractions of each category of cells during the control session (n = 365 cells from 6 mice). (E) Box plots of Ca^2+^ event rates of Chamber A-ON cells (A(E)-ON, n = 25 cells), Chamber B-ON cells (B(E)-on, n = 15 cells), Chamber A and B-ON cells (A(E)B(E)-ON, n = 8 cells) and other cells whose activity was not (anti-)correlated with investigation of chambers (Other, n = 311 cells) during the periods when the subject mice investigated Chamber A (A), Chamber B (B) or otherwise (C) in control sessions. Whiskers represent 10–90 percentile and red dots represent outliers. Cell categories whose fractions are larger than 1% are shown. ***a, P = 0.0001; ***b, P = 0.0004; **c, P = 0.0081; **d, P = 0.0081; ns^e^, P > 0.99; Friedman test with Dunn’s multiple comparisons test. (F) A pie chart showing the fractions of each category of cells in the first interaction session (n = 365 cells from 6 mice). (G) Box plots of Ca^2+^ event rates of Chamber A-ON cells (A(O)-ON, n = 8 cells), Chamber A-OFF cells (A(O)-OFF, n = 5 cells) Chamber B-ON cells (B(S)-ON, n = 59 cells), Chamber B-OFF cells (B(S)-OFF, n = 15 cells) Chamber A and B-ON cells (A(O)B(S)-ON, n = 4 cells) and other cells (n = 272 cells) in the first interaction session. **a, P = 0.0081; *b, P = 0.034; ***c, P < 0.0001; ns^d^, P = 0.085; ***e, P = 0.0004; *f, P = 0.04; ns^g^, P > 0.99; Friedman test with Dunn’s multiple comparisons test. (H) A pie chart showing the fractions of each category of cells in the second interaction session (n = 365 cells from 6 mice). (I) Box plots of Ca^2+^ event rates of Chamber A-ON cells (A(S)-ON, n = 41 cells), Chamber A-OFF cells (A(S)-OFF, n = 12 cells), Chamber B-ON cells (B(O)-ON, n = 28 cells), Chamber A and B-ON cells (A(S)B(O)-ON, n = 6 cells) and other cells (n = 273 cells) in the second interaction sessions. ***a, P < 0.0001; ***b, P = 0.0007; ***c, P < 0.0001; **d, P = 0.0045; ns^e^, P = 0.74; Friedman test with Dunn’s multiple comparisons test. (J) Changes in the fractions of A-ON, A-OFF, B-ON and B-OFF cells across sessions.

Subject mice spent comparable extent of time nose-poking to empty chambers A and B during control experiments (Fig. 3B). During the first interaction session, however, the amount of time nose-poking to Chamber B that housed a stranger mouse substantially increased and was much more than that to Chamber A that contained a novel object (Fig. 3B). During the second interaction session, the subject mice again spent significantly more time nose-poking to chamber A with a stranger mouse than to chamber B with a novel object, although the difference in time spent investigating the two chambers appeared less prominent than during the first session. The subject mice spent 82.9 ± 4.9 % and 84.5 ± 4.5% of total time immobile in the first and second interaction sessions, respectively. In addition, mice were almost completely stationary while they were nose-poking to each of the test chambers (93.0 ± 4.5 % and 97.5 ± 1.1 %, Chamber A and Chamber B during the first interaction session, respectively; 96.7 ± 3.8 % and 95.1 ± 3.4 %, Chamber A and Chamber B in the second interaction sessions, respectively). Overall, these results demonstrate that a subject mouse investigates the chamber that contains a stranger mouse more than that of the novel object, irrespective of the chamber’s location.

We then sought neuronal activity correlated or anti-correlated with the investigation of chamber A and/or B (Fig. 3C). Of 365 cells from 6 mice, 6.9% (25 cells) and 4.1% (15 cells) of total cells showed activity significantly correlated with the period of nose-poking to empty chamber A and chamber B, respectively, during control experiments (Fig. 3D). These two types of cells showed higher calcium event rates during investigation of chambers that their activity was correlated with compared to the other chamber (Fig. 3E). We thus categorized these cell types as “A(E)-ON cells” and “B(E)-ON cells”, in which the labels represents the chamber they responded to, followed by the content of the chamber in parenthesis (i.e. “E” for empty, “S” for stranger and “O” for object) and the type of response (“ON” for activation and “OFF” for suppression). Similarly, we found that 0.5% (2 cells) and 0.8% (3 cells) of total cells showed activity significantly anti-correlated with chamber A (A(E)-OFF cells) and B (B(E)-OFF cells), respectively, and 2.2% (8 cells) and 0.3% (1 cells) of total cells exhibited activity significantly correlated (A(E)B(E)-ON cells) and anti-correlated (A(E)B(E)-OFF cells) with investigation of chamber A as well as chamber B (Fig. 3D, 3E, and 3J). These findings indicate that AI contains a range of small fractions of neurons that exhibit diverse responses during investigation of the chambers.

In the first interaction session, the fractions of cells that exhibited activity significantly correlated (“B(S)-ON cells”) and anti-correlated (“B(S)-OFF cells”) with social investigation towards chamber B increased to 16.2% (58 cells) and 4.1% (15 cells) of total cells, whereas those that showed activity correlated and anti-correlated with non-social investigation towards chamber A remained low (“A(O)-ON cells”, 2.2% [8 cells]; “A(O)-OFF cells”, 1.4% [5 cells]; Fig. 3F, 3G and 3J). During the second interaction session, where the chambers that contained a stranger mouse and a novel object were exchanged, the fractions of cells whose activity was significantly correlated (“A(S)-ON cells”) and anti-correlated (“A(S)-OFF cells”) with nose-poking to chamber A increased to 11.2% (41 cells) and 3.3% (12 cells) of total cells, whereas those correlated and anti-correlated with nose-poking to chamber B decreased to 7.7% (28 cells. “B(O)-ON cells”) and 0.5% (2 cells, “B(O)-OFF cells”) of total cells (Fig. 3H, 3I and 3J). These results indicate that social stimuli activate a larger fraction of AI neurons compared to non-social stimuli. Moreover, this increase occurs independent of the locations at which the subjects investigate the social targets and accompanies parallel increases of cells that reduce their activity during social investigation.

We further examined whether the preference of activity of the cells of each category persisted across sessions. More specifically, we focused on the cells of the following categories (Fig. 4A): (1) cells that were consistently activated or suppressed during investigation of chambers with stranger mice or with novel objects (here termed “Social cells” and “Object cells” for convenience), (2) cells that were activated or suppressed during investigation of chamber A and chamber B only when they were not empty (“Conditional chamber A cells” and “Conditional chamber B cells”), and (3) cells that were activated or suppressed during investigation of chamber A and chamber B regardless of whether the chambers were empty or not (“Chamber A cells” and “Chamber B cells”). We found that 4.4% (16 cells) of total cells were Social cells (Fig. 4B and 4C and Supplementary Fig. 3). In addition, 1.4% (5 cells), 3.0% (11 cells) and 1.1% (4 cells) of total cells were Object cells, Conditional chamber B cells and Conditional chamber A cells, respectively (Fig. 4B and 4C). However, we found no Chamber A cells and only 0.8% (3 cells) of total cells were Chamber B cells. These results indicate that the fraction of cells encoding social exploration is larger than that of cells encoding exploration of non-social stimuli. Furthermore, neurons encoding spatial information *per se* are rare in AI.

**Figure 4.**
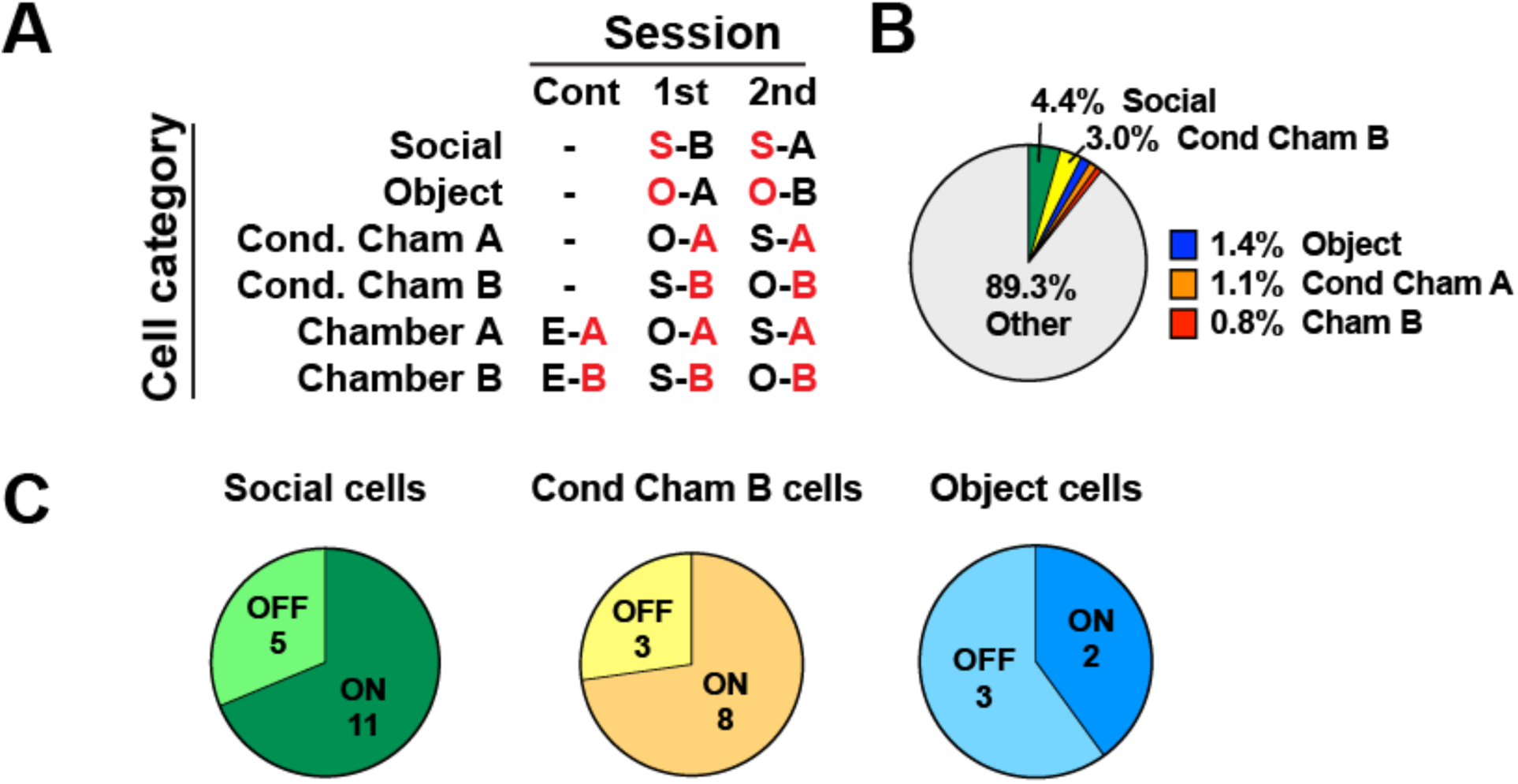
Social cells and other cell categories in LC experiments. (A) Definition of each category of cells and their preference for stimulus-chamber relationship. A and B represent chamber A and chamber B, respectively, and S, O and E represent stranger, object, and empty, respectively. The chamber and its content are connected by a hyphen to represent stimulus-chamber relationship. The features each cell category preferentially responds to are shown in red. (B) A pie chart that shows the fractions of each cell category within the total cell population (n = 365 cells from 6 mice). (C) Pie charts showing the fractions of “ON” cells and “OFF” cells within total Social cells (left), total Conditional chamber B cells (middle) and total Object cells (right). Numbers represent absolute cell numbers.

Finally, we examined whether insular social cells identified in the two different tasks formed a shared neuronal subset. Among 61 B(S)-ON cells and 17 B(S)-OFF cells identified during the first interaction session of the LC experiment, 10 B(S)-ON cells (16.4%, found in four out of six subject mice) and 1 B(S)-OFF cells (5.9%, found in one out of six mice) were also Social-ON cells and Social-OFF cells, respectively, in HC experiments (Supplementary Fig. 4). Moreover, 27.3% of the Social-ON cells that were consistently activated across multiple sessions in LC experiments (three out of 11 cells, found in three out of six mice) were also Social-ON cells in HC experiments, although none of five Social-OFF cells overlapped with those in HC experiments. These shared social cells observed in multiple subjects suggests the existence of a core subset of social cells acting across different social situations, whereas their unshared subpopulations operate in a task- and/or context-specific manner.

## Discussion

A previous whole-brain mapping study demonstrated that c-fos promoter-driven GFP-expressing neurons increased in many brain areas including the AI after social encounter (Kim et al., 2015). Our calcium imaging study significantly expands this observation and advances our understanding of how AI neurons encode social behavior at a single cell level by providing new findings that are difficult to observe with immediate early gene mapping. One such finding is that a small subset of Social-OFF cells emerged together with Social-ON cells during social exploration. The Social-ON and Social-OFF ensembles can potentially increase signal-to-noise ratio of information coding at a population level and can separately route opposing information via segregated projections to distinct downstream neurons. A homeostatic compensatory mechanism to balance the circuit activity may underlie the parallel emergence of Social-ON and Social-OFF cells. Since inhibition plays a fundamental role in shaping responses of cortical neurons, it will be important to examine the role of inhibition in emergence and lifetime development of these neuronal subsets in AI (Gogolla et al., 2014). Similar ON and OFF neural ensembles that code social exploration are recently reported in the prelimbic area (PL) of mPFC (Liang et al., 2018). Anatomical studies demonstrate that AI directly projects to PL but not vice versa (Allen et al., 1991; Shi and Cassell, 1998), raising the possibility that Social-ON and Social-OFF cells in AI may provide feedforward inputs to the ON and OFF ensembles in the PL.

Another remarkable finding is that activities of most Social-ON cells have their own preference for the target of physical contact and behavioral states. The multisensory nature of AI is well-suited for shaping such multidimensional representations of social exploration. A social target provides sensory cues of multiple modalities such as olfaction, audition, vision and touch, and IC receives sensory inputs of these modalities via thalamic and horizontal cortical afferents (Gogolla, 2017; Allen et al., 1991; Shi and Cassell., 1998). The higher fraction of Social cells observed in HC experiments compared to LC experiments could be explained by the difference in direct contact to social targets, which not only includes tactile stimulus but also activates accessory olfactory bulb when combined with pheromonal stimulation (Luo et al., 2003). Unique combinations of different social cues at each target of physical contact may most likely drive a subset of Social-ON cells preferentially over others. Regarding behavioral state preference, we found “Move cells” and “Stat cells” which were preferentially activated during particular behavioral states irrespective of social interaction. This implies that activities of Social-move cells and Social-stat cells could be shaped by convergence of such behavioral state signals onto subsets of Social cells

In LC experiments, social stimulus markedly increased the number of AI neurons that changed their activity upon nose-poking to that chamber. Although neurons consistently encoding the chamber location were hardly found, presentation of a stranger mouse or a novel object increased the neurons that were responsive to these chambers. These findings highlight an important feature of AI neurons in that they encode exploration of salient stimuli rather than spatial information itself. The expanded fraction of cells that encoded social investigation in prior sessions may have some influence on population coding in later sessions, as we observed more cells that consistently encoded the location of the chamber with a social stimulus in the first session (Conditional chamber B cells) compared to those that encoded the location of the other chamber (Conditional chamber A cells). It would be interesting in future research to test whether these cells encode a form of memory regarding the location of social stimuli or experience of social interaction.

In contrast to a wealth of findings on the social function of IC in humans, much less is known about its cellular and circuit foundations. Human neuroimaging studies have also implicated IC in salience detection and self-awareness (Craig, 2009; Uddin, 2014). Consistent with this view, our findings imply that a primary function of insular social cells is to encode the ongoing status of social interaction. The finding that only a minor fraction of AI neurons exhibited social interaction-related activity likely reflects multifunctionality of the AI and suggests that they may interact with other ensembles that are engaged in different aspects of cognition and behavior (Jennings et al., 2019). Finally, the structure and function of IC are known to be altered in various brain disorders including schizophrenia and ASD. Our study thus provides a cellular basis of insular social function on which future research using mouse models of neuropsychiatric and neurodevelopmental disorders will be grounded (Nakai et al., 2018; Takumi et al., 2019).

## Methods

### Ethics statement

All experimental procedures were conducted in accordance with institutional guidelines and protocols approved by the RIKEN Animal Experiments Committee.

### Mice

Male C57BL/6J mice (8–12 weeks-old, Japan SLC, Inc., Hamamatsu, Japan) were used. Mice were maintained on a 12 h-12 h light-dark cycle with *ad libitum* access to food and water. All surgical manipulations and behavioral tests were conducted during the dark phase of the cycle.

### Surgery

Each mouse underwent two separate surgical procedures. In the first surgery, the mouse was subjected to microinjection of an adeno-associated viral (AAV) vector that drove neuronal expression of a genetically-encoded calcium indicator and insertion of a gradient refractive index (GRIN) lens into the insular cortex (AI). In the second surgery, a baseplate for the miniaturized head-mounted microscope (nVista, Inscopix, Palo Alto, CA) was affixed onto the skull.

The AAV vector injection and GRIN lens insertion were conducted on the same day. Mice anaesthetized with 2% isoflurane in oxygen at a flow rate of 0.5L/min were placed in a stereotactic setup (Kopf Instruments, Tujunga, CA). Body temperature was maintained at 37°C with a heating pad (40-90-2-05, FHC, Bowdoin, ME). Ophthalmic ointment (Sato Pharmaceuticals, Tokyo, Japan) was applied to the eyes to prevent drying. A piece of scalp was removed and a small craniotomy was performed on the skull above the right AI using a high-speed rotary micro drill (OmniDrill 35, World Precision Instruments, Sarasota, EL). A 35-gauge needle attached to a microsyringe (World Precision Instruments) was targeted to the right AI (1.94 mm anterior, 2.2 mm lateral and 3.45 mm in depth relative to bregma), and 500 nl of virus solution (AAV5-CaMKII-GCaMP6f-WPRE-SV40, 1.0×10^14^ / mL, UPenn Vector Core) was microinjected at a rate of 100 nl/min. The needle was left undisturbed for additional 10 min to avoid backflow. The needle was then slowly withdrawn and the injected surface was washed with saline. Following the microinjection, a GRIN lens (0.5 mm diameter, 4.0 mm length, GLP-0540, Inscopix, Palo Alto, CA) attached to a lens implant kit (ProView, Inscopix, Palo Alto, CA) was slowly inserted stereotaxically to a position slightly dorsal to the AAV injection site (1.94 mm anterior, 2.2 mm lateral and 3.25 mm in depth relative to bregma). The lens was then affixed to the skull with dental cement and the top of the lens was covered with a silicon mold (Kwik-Cast, World Precision Instruments, Sarasota, EL). Mice were singly housed after fully recovering from the surgery.

Four weeks after the GRIN lens implantation, mice were anaesthetized again with isoflurane and the silicon mold over the tip of GRIN lens was carefully removed. The baseplate (Inscopix) attached to a miniaturized head-mounted microscope was positioned above the implanted lens using an adjustable gripper (Inscopix). The microscope and baseplate were then lowered towards the top of the lens until the field of view was in focus. After confirmation of GCaMP fluorescence signals, the baseplate was affixed to the skull using dental cement. The baseplate was covered with a baseplate cover (Inscopix) after the microscope was detached from the baseplate.

### Social behavior tests

We used two different behavioral assays, the home-cage (HC) experiment and the linear chamber (LC) experiment, for social behavior tests. Subject mice were singly housed in their HCs for at least 4 weeks before the exposure to social stimuli. Behavioral schedule consisted of at least two days of habituation and a day of testing for HC experiments, followed by at least two days of habituations and a day of testing for LC experiments. A total of nine mice were used for these experiments. Data from all nine mice were analyzed in HC experiments, and data from three mice were excluded in the analysis of LC experiments because imaging data from these mice contained less than 30 cells. The behavior was recorded using a video camera (logicool) throughout the test in both paradigms. Different stranger mice were used in HC experiments and LC experiments.

In the HC experiment, a subject mouse with a microscope attached to its head underwent habituation sessions in which the mouse was allowed to move freely in its HC (30 cm long, 18 cm wide and 12 cm high) for at least 20 min a day. On the day of imaging, a set of tests that consisted of a control session and an interaction session were performed as follows (Fig. 1A). First, a novel miniature object (nonsocial target) and a microscope-attached subject mouse were placed together in its HC, and the mouse was allowed to move freely in this environment for 4 min. Immediately after this control session, the object was replaced with a stranger mouse (social target) and the subject mouse was allowed to freely explore for an additional 4 min. The subject mouse remained in the test environment during the brief interval between the two sessions.

In the LC experiment, an acrylic LC that consisted of a center chamber (40 cm long, 10 cm wide and 21 cm high) flanked by two test chambers (chambers A and B; 10 cm long, 10 cm wide and 21 cm high each) was used (Fig. 3A; Lee et al., 2016). The bottom 10 cm of the walls that divided the center chamber and the test chambers had 1-cm diameter holes drilled to allow subject mice to interact with the targets by nose poking. Each test chamber contained a stranger mouse (social stimulus), a miniature object (nonsocial stimulus) or nothing (no stimulus). At least 2 days prior to imaging experiments, a subject mouse with a microscope attached to its head was allowed to move freely in the center chamber during a 20-min habituation session a day for two days. The test consisted of three consecutive sessions (Fig. 3A). In the first session, a subject mouse placed in the center chamber was allowed to move freely for 4 min while no stimuli were presented in the two test chambers (control session). In the following session, a novel miniature object and a stranger mouse were placed in chamber A and chamber B, respectively, and the subject mouse was allowed to freely explore in the center chamber for an additional 4 min (first interaction session). In the last session, the positions of the stranger mouse and the object were swapped and the subject mouse was allowed to explore freely for another 4 min (second interaction session).

### Behavioral data analysis

The behavior of subject mice was manually classified by visual inspection of the videos after the experiments. In HC experiments, we defined the periods of social interaction as those during which the subject mouse actively explored the social target by nasal contact, and these periods were further subclassified into the periods of contact to the nose, body and anogenital area of the stranger mouse. The remaining periods were categorized as non-social periods. Within the non-social periods, the periods during which the subject mouse touched the wall of the HC and those during which the subject mouse was passively touched by the stranger mouse were documented. For analysis of behavioral states, the periods during which the subject mouse changed its body and hindlimb positions were defined as moving periods, and the remaining periods of immobility were classified as stationary periods. In LC experiments, we defined the periods of investigation of the test chamber as nose-poking by the subject mouse to the wall that divided the center chamber and the test chamber.

### Ca^2+^ imaging of AI neurons in freely-moving mice

Activity of GCaMP6f-labeled AI neurons during social behavior tests was imaged using a miniaturized head-mounted microscope and GRIN lens-mediated microendoscopy (Supplementary Fig. 1A). Before the experiments, mice were lightly anesthetized with 2 % isoflurane and the baseplate cover was removed from the baseplate, to which a miniaturized head-mounted microscope was subsequently attached. Mice were then recovered from anesthesia for at least 20 min before beginning the experiments.

Ca^2+^ imaging videos were recorded during social behavior tests using nVista acquisition software (version 1.2.0; Inscopix, Palo Alto, CA) with a resolution of 1440 × 1080 pixels at a rate of 15 frames per second. The LED power of the microscope was set between 30–50% (0.36–0.6 mW). Ca^2+^ transients were observed in many neurons within the field of view (FOV) while mice moved freely in their environments (Supplementary Fig. 1B, 53.6 ± 24.4 cells/FOV, mean SD, n = 9 mice), and post-hoc confirmation of the lens positions verified that GRIN lenses were successfully targeted to AI in all nine cases (Supplementary Fig. 1C).

### Imaging data analysis

Post-acquisition processing of Ca^2+^ imaging videos was performed using the Mosaic software (version 1.0.5b; Inscopix). Videos were spatially downsampled by a factor of 4. Motion correction was performed by shifting each frame to a single reference frame so that high contrast features within each frame were aligned to the corresponding features in the reference frame. Normalized fluorescence changes were then visualized in a ΔF/F_0_ video, in which a minimum z-projection image of the entire movie (F_0_) was subtracted from each frame (F) and the resultant F-F_0_ movie was normalized to F_0_. Cells in the ΔF/F_0_ video were then identified using an independent-component-analysis- and principal-component-analyses-based automated cell-segmentation algorithm in Mosaic, followed by human visual verification. Ca^2+^ transient events were detected using a Ca^2+^ event detection algorithm in Mosaic (parameters, tau = 200ms, Ca^2+^ transient event minimum size = 6 median average deviation) by finding large peaks of fluorescence changes with fast rise times and exponential decay.

Neurons that exhibited activity correlated and anti-correlated with social behavior were searched by statistical testing of cosine similarity between the binary vector A that represented the occurrence of neuronal activity event by 1 and the binary vector B that represented the period of social interaction by 1. Cosine similarity between these two vectors was calculated as follows:

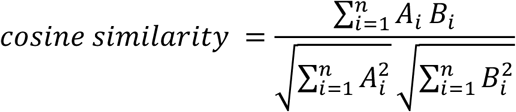

We compared this value to a distribution of cosine similarity calculated using 1000 randomly permuted data derived from the same cell. The permutation was conducted by rotating the activity event time series by a random amount relative to the time series of behavioral data. Cells were considered “ON cells” and “OFF cells” if their cosine similarity in the real data was greater than the 95th percentile and smaller than the 5th percentile of the values obtained from the randomly permuted data, respectively. Identification of neurons whose activity was correlated with other types of behavior was conducted similarly by using the binary vector that represented the occurrence of the behavior of interest by 1.

Ca^2+^ event rate of each neuron during a particular behavior was calculated as the number of Ca^2+^ transient events during the periods of the behavior of interest divided by the total length of the periods. To quantitatively estimate the preference of activity of each neuron for social interaction, we calculated the social preference index as (S-N)/(S+N), where S and N are Ca^2+^ event rates of the neuron during social interaction and non-social periods, respectively. The social preference index ranges from −1 to 1, where a positive value indicates a preference for social interaction and a negative value indicates a preference toward non-social periods. Calculation of nearest neighbor distances (NNDs) between Social-ON cells was conducted by averaging NNDs across all Social-ON cells within the map. To test whether the anatomical distribution of Social-ON cells was random, the same numbers of labels of Social-ON cells as the real data were randomly shuffled 1000 times and an average of NNDs calculated from the shuffled dataset was compared to the values obtained from the real data. Since microendoscopic images obtained through a GRIN lens are subjected to image distortion at periphery, we expressed NNDs in arbitrary relative distances (i.e. in pixels) rather than in absolute distances. Identification of Social cells that were common between HC experiments and LC experiments were conducted by visual comparisons of the Social cell maps obtained in these two experiments.

### Histology

Mice deeply anesthetized with isoflurane were transcardially perfused with phosphate-buffered saline (PBS) followed by 4% paraformaldehyde (PFA) in PBS. Brains were dissected out and post-fixed in 4% PFA overnight. 50 µm-thick coronal sections were cut on a vibratome (Leica VT1200S). Fluorescence images were acquired using a Keyence BZ9000 epifluorecence microscope (Keyence, Osaka, Japan).

### Statistics

Data are represented as mean ± SEM. Statistical tests were performed using GraphPad Prism version 8 (GraphPad Software, Inc., La Jolla, CA). All tests were two-sided, and statistical significance was defined as P < 0.05. Exact *P*-values are shown unless *P* < 0.0001.

## Author contributions

IM, NN and TT conceived the study. IM, NK and ETNO conducted the experiments. IM and MS analyzed the data. IM, MS and TT wrote the paper with input from JN and TK.

## Acknowledgement

This work was supported by KAKENHI (16H06316, 16H06463) from Japan Society for the Promotion of Science (JSPS) and Ministry of Education, Culture, Sports, Science, and Technology, Intramural Research Grant for Neurological and Psychiatric Disorders of NCNP, the Takeda Science Foundation and Smoking Research Foundation to TT, KAKENHI (17H05985, 19H04942) to M.S., and KAKENHI (15H05723) to J.N.

---

**Supplementary Figure 1.**
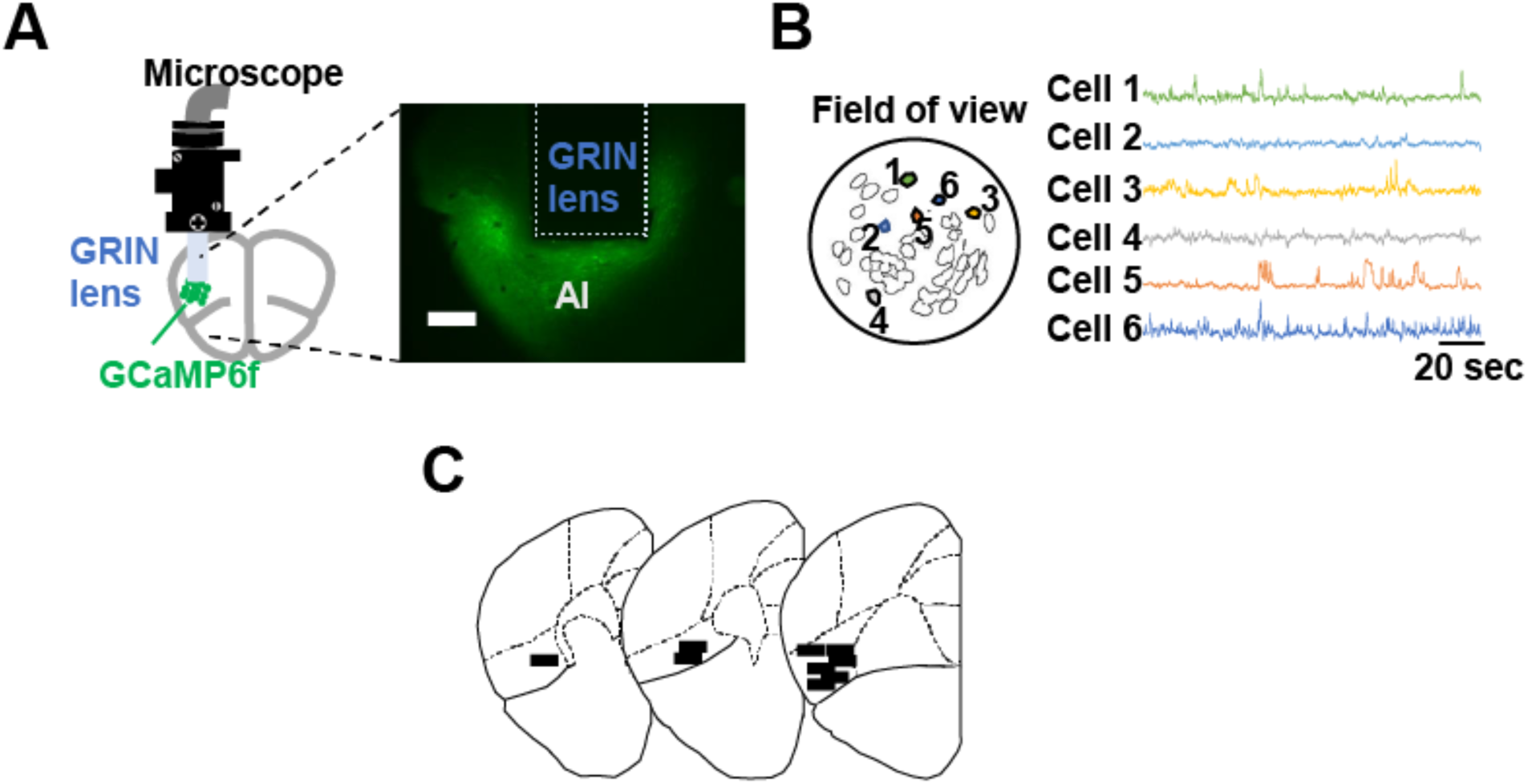
Microendoscopic calcium imaging of AI neurons in freely-moving mice. (A) A schematic of microendoscopic imaging (left). AI neurons virally transfected with GCaMP6f were imaged using a miniaturized head-mounted microscope through a chronically implanted GRIN lens. A representative fluorescence image of a coronal section shows the trace of a GRIN lens implanted close to GCaMP6-expressing AI neurons (right). Scale bar = 200 µm. (B) A representative field of view showing contours of a population of imaged AI neurons (left). Six example neurons are numbered and their time-varying GCaMP6f fluorescence changes are shown on the right side. (C) Reconstructed positions of the implanted GRIN lenses in AI (n = 9 mice). Approximate centers of the lenses along the anteroposterior axis are shown by black horizontal lines in drawings of representative coronal sections obtained from the Allen Brain Atlas.

**Supplementary Figure 2.**
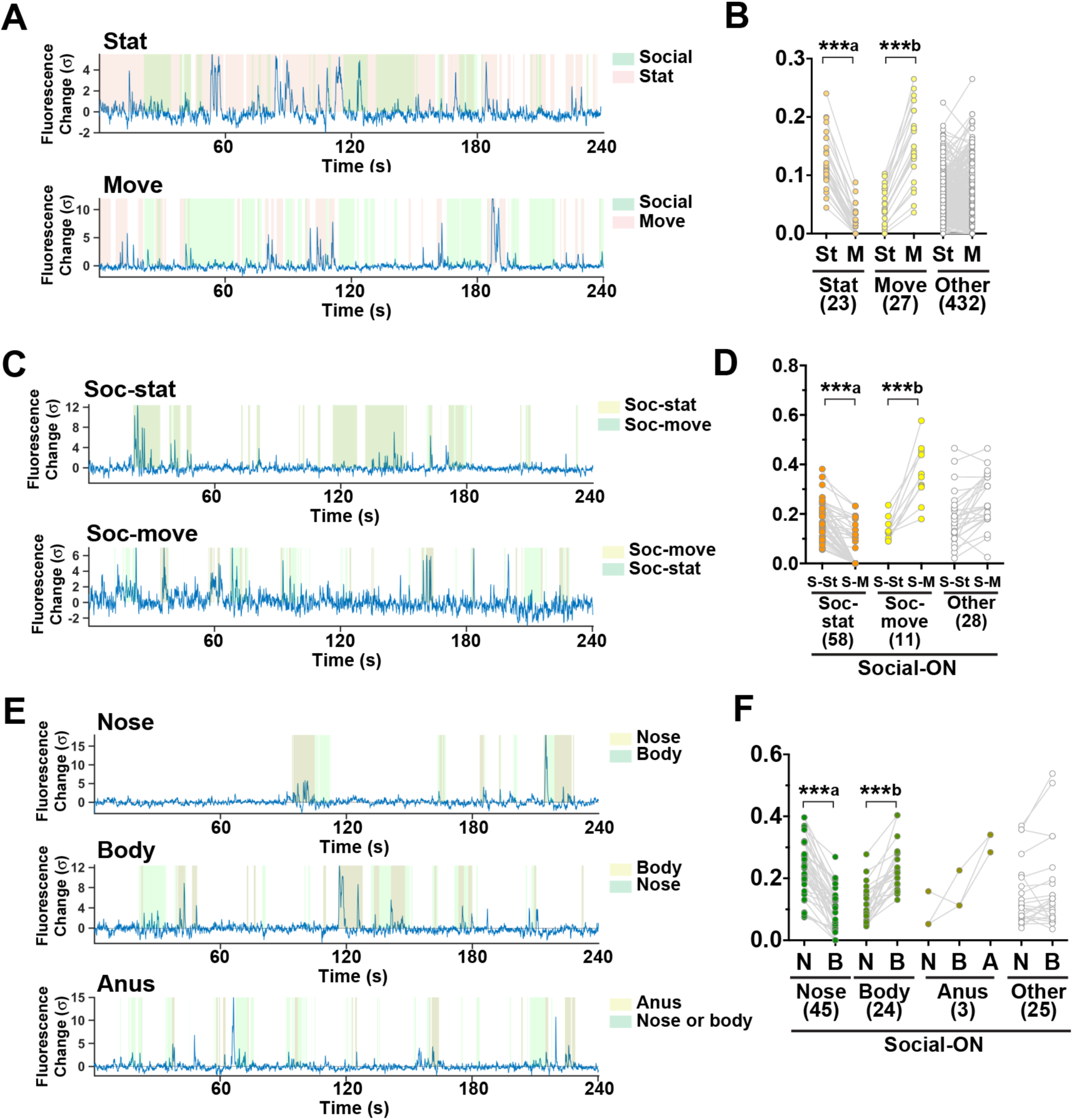
Ca^2+^ event rates of each cell type during social behavior in HC experiments. (A) GCaMP6f fluorescence changes of a Stat cell (top) and a Move cell (bottom) during 4-min interaction sessions. (B) Ca^2+^ event rates of Stat cells, move cells and other cells during the stationary (St) and moving (M) periods. Numbers of each cell type are shown in parentheses. Event rates of the same cells are connected by gray lines. The same convention applies to D and F. ***a, P < 0.0001, W_(23)_ = −276; ***b, P < 0.0001 W_(27)_ = 378; Wilcoxon matched-pairs sign rank test. (C) GCaMP6f fluorescence change of a Social-stat cell (top, Soc-stat) and a Social-move cell (bottom, Soc-move). (D) Ca^2+^ event rates of Soc-stat cells, Soc-move cells and other Social-ON cells (other) during social interaction with (S-M) and without (S-St) movement. ***a, P < 0.0001, W_(58)_ = 1695; ***b, P = 0.001, W_(11)_ = −66; Wilcoxon matched-pairs sign rank test. (E) GCaMP6f fluorescence change of nose (top), body (middle), and anus (bottom) subtypes of Social-ON cells. (F) Ca^2+^ event rates of nose, body, anus and other subtypes of Social-ON cells during social interaction with contact with nose (N), body (B) and anus (A). Since the fraction of time spent contacting anus was low, only event rates during contact with nose and body are shown for nose, body and other cell subtypes. ***a, P < 0.0001, W_(45)_ = −1035; ***b, P < 0.0001, W_(24)_ = 300; Wilcoxon matched-pairs sign rank test.

**Supplementary Figure 3.**
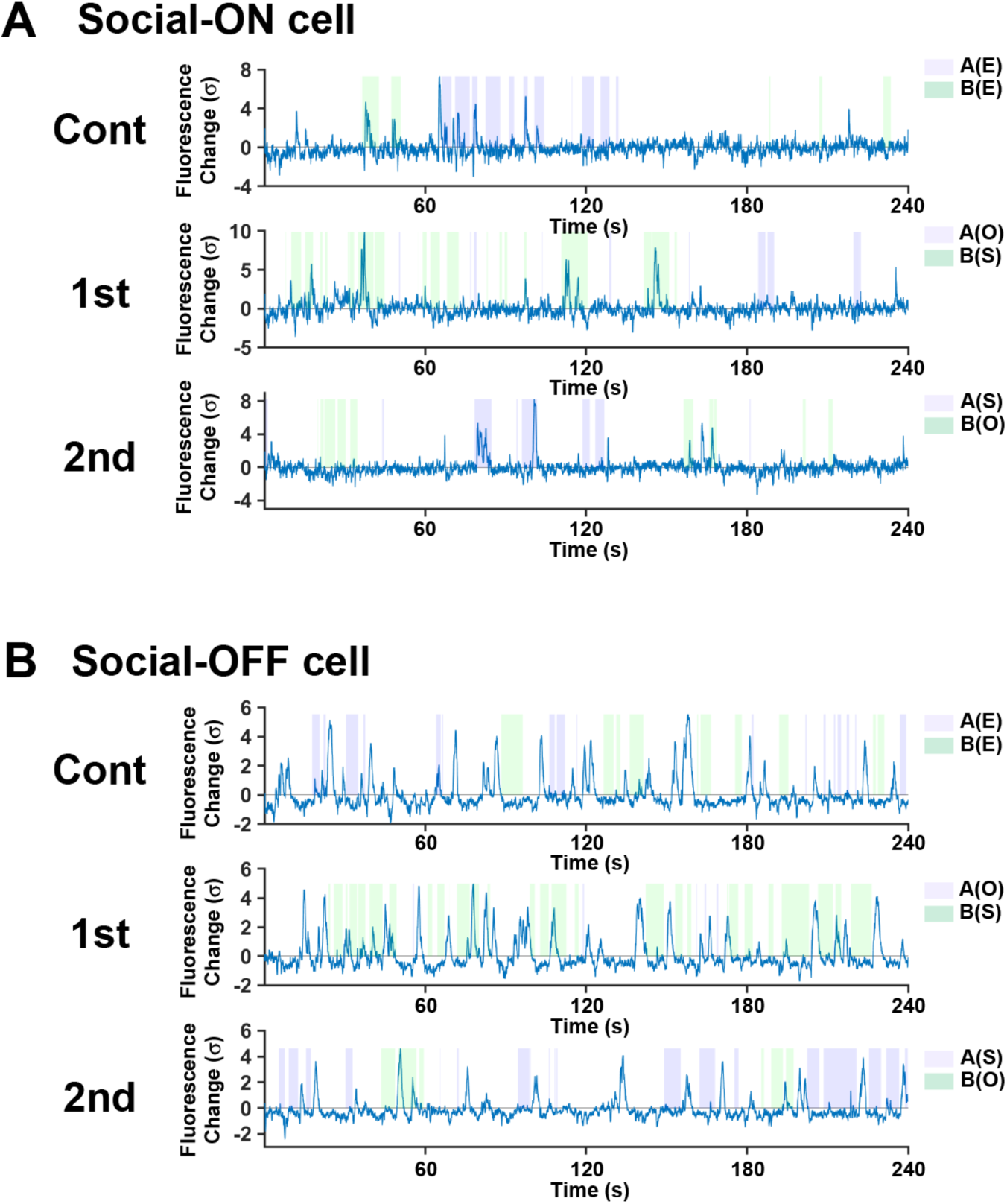
Social cells consistent across multiple LC sessions. (A). GCaMP6f fluorescence change of a Social-ON cell during control (top, Cont), first interaction (middle, 1st) and second interaction sessions (bottom, 2nd) of LC experiments. (B) GCaMP6f fluorescence change of a Social-OFF cell during control (top), first interaction (middle) and second interaction sessions (bottom) of LC experiments.

**Supplementary Figure 4.**
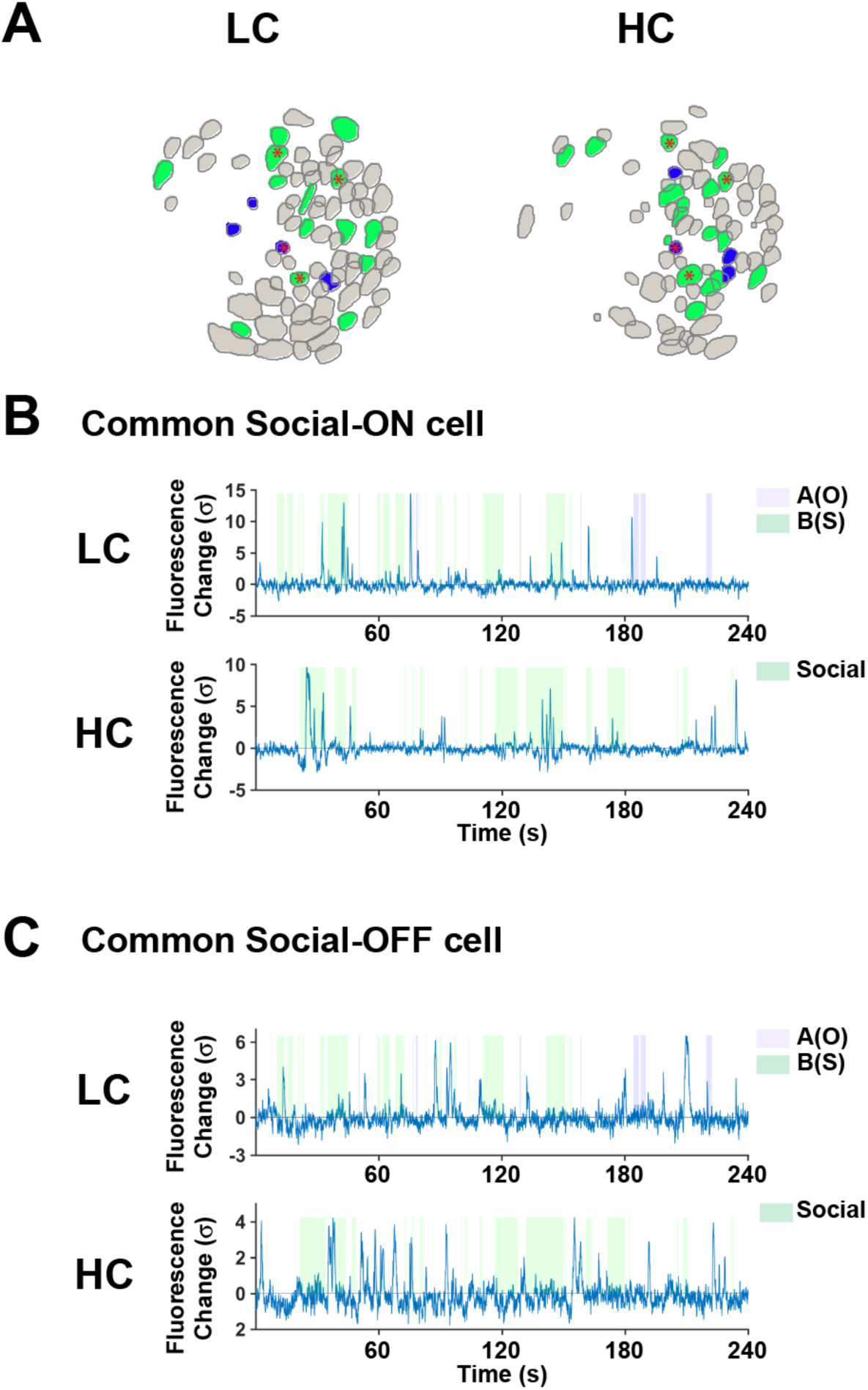
Social cells common across different tests in AI. (A) Example Social cell maps of LC experiments (left) and HC experiments (right) imaged in the same animal. Social-ON cells and Social-OFF cells are shown in green and blue, respectively. Social cells that are common to the two paradigms (common Social cells) are indicated by red asterisks. (B) GCaMP6f fluorescence change of a common Social-ON cell during the first interaction session of LC experiment (top) and HC experiment. (C) GCaMP6f fluorescence change of a common Social-OFF cell during a first interaction session of LC experiments (top) and a HC experiment.

